# Activity-based tyrosine phosphatomics using F_2_Pmp probes

**DOI:** 10.1101/2023.03.20.533451

**Authors:** Tomoya Niinae, Yasushi Ishihama

## Abstract

We showed that the F_2_Pmp probe binds to PTP in a sequence-dependent manner. In addition, this study is the first successful example of comprehensive enrichment of classical PTP at the protein level. Furthermore, we found that F_2_Pmp probe can enrich PTPs in a PTP activity dependent manner. Using the F_2_Pmp probe, the fluctuation of PTPN1 and PTPN2 activities were revealed. This enrichment approach using the F_2_Pmp probe could be a generic tool for activity-based tyrosine phosphatomics.

## INTRODUCTION

Tyrosine phosphorylation on proteins is reversibly regulated by protein tyrosine phosphatases (PTPs) and protein tyrosine kinases in living cells, and is closely related to various diseases including cancer^1,2^. More than 100 PTPs are known to exist in the human genome, among which class I PTPs, which have the catalytic cysteine residue in the active site, constitute the largest group^3^. The class I PTPs are divided into 63 dual-specificity PTPs (DUSPs), which target serine-threonine as well as tyrosine as substrates, and 37 tyrosine-specific classical PTPs. In addition, the classical PTPs are further divided into 20 receptor-type PTPs and 17 non-transmembrane PTPs.

It is important to have a general method for profiling the expression and activity of enzymes. As for the enzymes involved in phosphorylation, comprehensive enrichment methods for kinases and serine/threonine phosphatases have been developed^4,5^. For the comprehensive analysis of kinases, the enrichment method using pan-kinase inhibitor immobilized resin as a probe (Kinobeads) is known ^6^. Klaeger et al. applied this method to quantify more than 200 kinases and identify targets for kinase inhibitors in clinical use^7^. In addition, Lyons et al. developed a serine threonine phosphatase enrichment method using a resin immobilized with microcystin LR, a pan-serine threonine phosphatase inhibitor^8^. For PTPs, Karisch et al. developed a method to enrich peptide fragments of PTPs after enzymatic digestion using antibodies against highly conserved sequences in the active site^9^. However, no general approach for proteome-wide enrichment of PTPs has yet been reported.

Several probes that bind to PTPs have been developed^10–15^. For example, Kumar et al. developed an α-Bromobenzylphosphonate (BBP) probe and Liu et al. produced phenyl vinyl sulfonate (PVSN) and phenyl vinyl sulfone (PVS) probes^10–12^. Kalesh et al. produced a peptide probe containing the unnatural amino acid 2-FMPT and showed that the probe binds to PTPs in a probe-sequence-dependent and PTP activity-dependent manner^14^. However, although these reports showed that the probes bind to some recombinant PTPs and endogenous PTPs in cell extracts, they did not show whether endogenous PTPome could be enriched from cell extracts. In addition, there are off-targets (bound proteins other than PTPs) for these probes, but these have not been investigated much^16^.

In this study, we focused on a non-hydrolyzable phosphotyrosine analogue, F_2_Pmp, which is not dephosphorylated by PTP unlike phosphorylated tyrosine, shows high affinity for PTP and has been used to detect PTP in living cells and cell extracts^17–19^. Recently, we developed a method for protein-level PTP enrichment using F_2_Pmp, and succeeded in F_2_Pmp-dependent enrichment of PTPN1, PTPN2, PTPN13, and SH2 domain-containing proteins (RASA1, PIK3R1, and PIK3R2)^20^.

In this study, we aimed to develop a comprehensive enrichment method for PTPs and conducted more detailed studies on the design of systematic probe sequences and the correlation between probe sequences and enriched PTP. In addition, the correlation between the F_2_Pmp probe enrichment and PTP activity was evaluated.

## EXPERIMENTAL SECTION

### Cell culture

HEK293T cells were provided by RIKEN BRC through the National BioResource Project of the MEXT/AMED, Japan. HeLa cells (HSRRB, Osaka, Japan) and HEK293T cells were cultured in DMEM (Fujifilm Wako, Osaka, Japan) containing 10% fetal bovine serum (Thermo Fisher Scientific, Waltham, MA) and 100 µg/mL penicillin/streptomycin (Fujifilm Wako) in an incubator at 37 °C under humidified 5 % CO2 in air. Cells were washed with ice-cold phosphate-buffered saline (PBS), harvested with ice-cold PBS and frozen by liquid nitrogen.

### EGF treatment

For dual labeling SILAC, we used either [12C6, 14N4]-Arg (+0) and [12C6, 14N2]- Lys (+0) for “light” labeling or [13C6, 15N4]-Arg (+10) and [13C6, 15N2]-Lys (+8) for “heavy” labeling. In each SILAC condition, DMEM deficient in Arg and Lys was supplemented with stable isotope-encoded Arg and Lys, 10% fetal bovine serum, 100 µg/mL penicillin/streptomycin, 1% L-glutamine (200mM in 0.85% NaCl) and 1% sodium pyruvate (100mM). 90% confluent HeLa cells were incubated in serum-starved DMEM (0.1 % FBS) for 24 h. Heavy-labeled HeLa cells were stimulated with 150 ng/ml EGF for 2, 10, or 20 min. After the treatments, cells were washed with ice-cold phosphate-buffered saline (PBS), harvested with ice-cold PBS and frozen by liquid nitrogen.

### Peptide synthesis

All peptides used in this study probes were designed as 11-mers bearing biotinylated PEG2 linker in the N terminus and an amide group in the C terminus and were synthesized using Fmoc-solid-phase peptide chemistry with Novasyn TGR resin (Merck Millipore) as described in the Novabiochem Innovations 01/11. A coupling system using 1-((Dimethylamino)(dimethyliminio)methyl)-1*H*-[1,2,3]triazolo[4,5- *b*]pyridine 3-oxide hexafluorophosphate/*N,N*-diisopropylethylamine was employed for the introduction of F_2_Pmp (PEPTIDE INSTITUTE, INC., Osaka, Japan) and a coupling system using 1-hydroxybenzotriazole/2-(1H-Benzotriazole-1-yl)-1,1,3,3- tetramethyluronium hexafluorophosphat/*N,N*-diisopropylethylamine was employed for the introduction of all other amino acids and a biotin PEG2 acid (Broad Pharm, San Diego, CA). After synthesis, the resulting protected peptide resin was treated with a TFA cocktail [TFA/H2O/m-cresol/thioanisole/ethanedithiol (80:5:5:5:5)] for 3 h. The crude product was purified by reverse-phase HPLC.

### Peptide pulldown (cell lysate)

Per a pulldown experiment, 18 uL streptavidin-conjugated agarose beads (Thermo Fisher Scientific) were saturated with 160 nmol of biotinylated peptides in the lysis buffer. Cells were lysed in lysis buffer (1% (v/v) Nonidet P-40, 150 mM NaCl, 25 mM Tris-HCl (pH 7.5), and protease inhibitors (Sigma-Aldrich, St. Louis, MO)), and centrifuged at 15,800 g, 4 °C for 30 min. The resulting supernatant was used for the experiment. Protein concentration was determined by means of BCA assay (Thermo Fisher Scientific). An equal amount of proteins (3 mg) was incubated with the respective immobilized peptide probes at 4 °C for 4 h. After 3 rounds of washing with lysis buffer followed by rinsing with 100 mM Tris-HCl (pH 9), bound proteins were eluted using phase-transfer surfactant (PTS) buffer^21^ (12 mM sodium N-lauroylsarcosinate (FUJIFILM Wako), 12 mM sodium deoxycholate (FUJIFILM Wako), 100 mM Tris-HCl (pH 9), 10 mM dithiothreitol) at 95 °C for 5 min. Eluted proteins were filtered at 0.45 um.

### Peptide pulldown (recombinant PTPs)

Per a pulldown experiment, 18 uL streptavidin-conjugated agarose beads (Thermo Fisher Scientific) were saturated with 160 nmol of biotinylated peptides in the lysis buffer. Peptide conjugated resin was incubated with a mixture of 11 recombinant PTP (abcam, Cambridge, UK) (each 1.5 pmol) in 300 uL lysis buffer at 4 °C for 4 h. After incubation, the supernatant was incubated with a fresh peptide conjugated resin in 300 ul lysis buffer at 4 °C for 4 h. Then, the resin of first incubation and second incubation were washed and bound proteins were eluted as described above.

### Protein digestion

Eluted proteins were digested with Lys-C (FUJIFILM Wako) and trypsin (Promega, Madison, WI) according to the PTS protocol^21^, followed by desalting on SDB-XC (GL Sciences, Tokyo, Japan) StageTips^22^.

### Orthovanadate treatment

HEK293T cell lysate was incubated with 1.5 mM orthovanadate at 4 °C for 1 h.

### H_2_O_2_ treatment

Recombinant PTPN1 (0.5 pmol) was incubated with 1 mM H_2_O_2_ in the lysis buffer at 25 °C for 5 min. After the reaction, the reaction buffer was exchanged to the lysis buffer five times by Amicon Ultra 10K (Merck Millipore). Then, the sample was splitted to two. One was used for peptide pulldown, and bound PTPN1 was eluted, digested, desalted by SDB-XC and analyzed by LC/MS/MS as described above. The other was digested, cleaned up by SCX, desalted by SDB-XC and measured by LC/MS/MS.

### NanoLC/MS/MS

Nano LC/MS/MS was performed by an Orbitrap Fusion Lumos (Thermo Fisher Scientific) or timsTOF Pro (Bruker), connected to a Ultimate 3000 RSLCnano pump (Thermo Fisher Scientific) and an HTC-PAL autosampler (CTC Analytics, Zwingen, Switzerland) equipped with a self-pulled analytical column (150 mm length × 100 μm i.d.) packed with ReproSil-Pur C18-AQ materials (3 μm or 1.9 μm) (Dr. Maisch GmbH, Ammerbuch, Germany). The mobile phases consisted of (A) 0.5% acetic acid and (B) 0.5% acetic acid and 80% ACN and a flow rate was set at 500 nL/min. The following gradient was 5-10% B in 5 min, 10-40% B in 60 min or 100 min, 40-99% B in 5 min.

For the pulldown experiment against cell extracts, Orbitrap Fusion Lumos was employed. FAIMS separations were performed at CV = −40, −60, −80. The MS scan was performed at Orbitrap with a range of m/z 300-1,500 and a resolution of 120,000. The MS/MS scan was performed at Iontrap with Scan Rate of Rapid. Auto gain control was 100% for both MS scan and MS/MS scan. HCD was 30.

For the pulldown experiment against recombinant PTPs, timsTOF Pro was employed. PASEF number was 10. Ramp time and Accumulation time was 100 ms. Scan range of 1/K0 was 0.6–1.5 Vs/cm^2^, and Scan range of *m*/*z* range was 100–1700 *m*/*z*. Target value for PASEF-MS/MS scan was 24,000.

### Database searching

For all experiments, the raw MS data files were analyzed by MaxQuant v1.6.17.0^23^. Peptides and proteins were identified by means of automated database searching using Andromeda against the human SwissProt Database (version 2020-08, 20,368 protein entries) with a precursor mass tolerance of 20 ppm for first search and 4.5 ppm for main search and a fragment ion mass tolerance of 20 ppm. The enzyme was set as Trypsin/P with two missed cleavage allowed. Cysteine carbamidomethylation was set as a fixed modification. Methionine oxidation and acetylation on the protein N-terminus were set as variable modifications.

For pulldown experiments, match between run was performed between each F_2_Pmp and Y probe pulldown with the same sequence.

### Data analysis for the pulldown using cell lysate

For label-free quantification of peptide pulldown samples, only proteins quantified in 2 out of the 3 replicates in at least one condition were used for further analysis, and missing values were imputed from a normal distribution of log2 intensity using a default setting in Perseus^24^.

For the SILAC-based protein quantification of peptide pulldown samples, normalized H/L ratios were used for the further analysis.

### Data analysis for the quantification of affinity

Only PTP quantified more than twice out of three replicates, either in the first or second pulldown, is employed in the following analysis, and missing values were imputed the smallest intensity among the other PTP in the same measurement. The calculation of the dissociation constant *K_d_* was performed as previously reported by Sharma et al.^25^ and the following equation was used.

In addition, PTP with *r* greater than 1 in any one of the three replicates was excluded from the calculation of *K_d_* values.

### PWM score

We employed peptides, which have value >2 in the peptide array experiment^26^, as the PTP substrates. The probability of observing residue in position from *in vitro* substrates for PTPs is computed as follows:

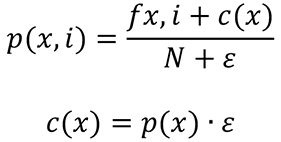

where fx,i is the frequency of observing residue x at position i and N is the total number of sequences. c(x) is a pseudo count function which is computed as the probability of observing residue x in the peptide used in the peptide array experiment, p(x), multiplied by E, defined as the square root of the total number of sequences used to train the PWM. This avoids infinite values when computing logarithms. Probabilities are then converted to weights as follows:

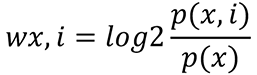

where p(x) *=* background probability of amino acid x; p(x,i) = corrected probability of amino acid in position; Wx,i *=* PWM value of amino acid in position. Given a sequence q of length I, a score λ is then computed by summing log_2_ weights:

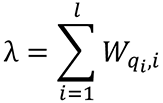

where qi is the ith residue of q. In this study, the score was computed using the flanking ±5 residues surrounding each phosphosite.

### Motif Enrichment

rmotifx (https://github.com/omarwagih/rmotifx)^27^ was used to identify motifs enriched. Motif enrichment was performed with a set of pY sites in PhosphoSitePlus (foreground) and human proteome (background) at minimum sequence ≧ 1,000 and p < 1e-6.

## RESULTS AND DISCUSSION

### Measurement of dissociation constants between F_2_Pmp probe and recombinant PTP

To investigate the relationship between more diverse probe sequences and PTPs, 17 F_2_Pmp peptide probes shown in Table 1 were synthesized (biotin was attached to the N-terminus via a PEG_2_ linker). Among them, ten probes are designed based on pY sites present in human cells^28^. Four probes are substitutions of amino acids in the −1 position of PDGFRB Y1021. In addition, DIDOI (Y1016) was selected because the pY peptide of the same sequence has been reported to bind to many PTPs^26^. Other probes were synthesized that consisted of F_2_Pmp only or F_2_Pmp and two residues. These probe sequences contained 8 of the top 10 specific motifs around the pY sites registered in PhosphoSitePlus, and the pY sites with these 8 motifs were approximately half of all pY sites registered in PhosphoSitePlus (Table S1, Figure S1A)^28^.

**Table 1.**
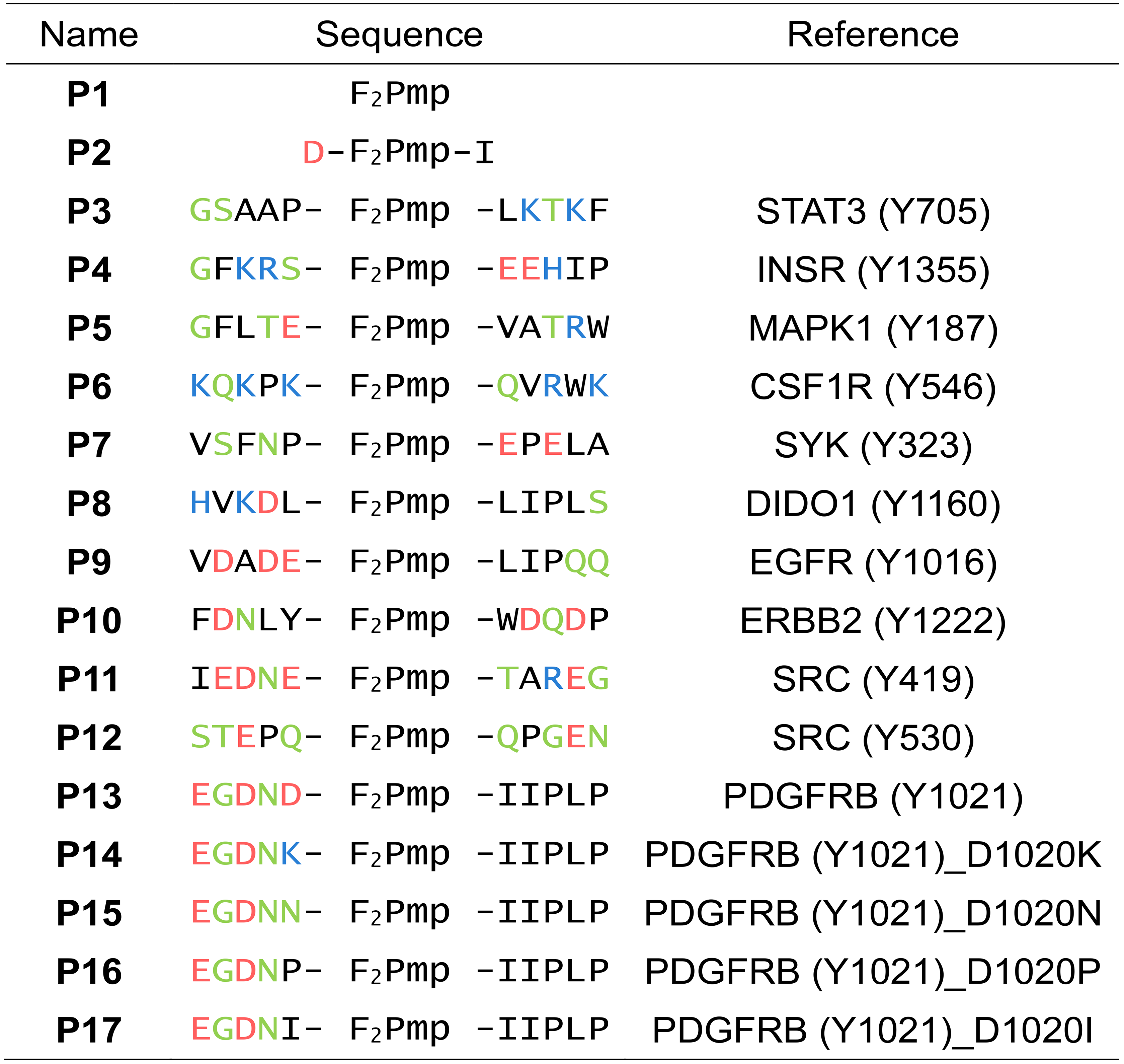
List of the used peptide probes. Each peptide comprises a biotinylated PEG_2_ linker in the N terminus.

To evaluate the binding between the F_2_Pmp peptide probe and PTPs quantitatively, we measured the dissociation constant *K_d_* between the F_2_Pmp probe and PTPs using a method developed by Sharma et al^25^. That is, the F_2_Pmp probe was conjugated to a streptavidin resin and then incubated with a mixture of 11 recombinant PTPs (10 classical PTPs and 1 DUSP). After the first incubation, the supernatant fraction was again incubated with a fresh F_2_Pmp probe conjugated resin, and the *K_d_* of the F_2_Pmp peptide-PTP complex was calculated from the change in the intensity of bound PTP in the two pulldowns. The *K_d_* value of many F_2_Pmp peptide probe/PTPN1 complexes was several hundred nM to several μM (Figure 1). This result is consistent with the previously reported dissociation constants between PTPN1 and F_2_Pmp probes^29^. In addition, when compared with the enrichment ratios using tyrosine probe as a control, a positive correlation was observed between the inverse logarithm of *K_d_* and the logarithm of F_2_Pmp /Y ratio, indicating that the binding of PTP to F_2_Pmp probe is F_2_Pmp dependent (Figure S1B).

**Figure 1.**
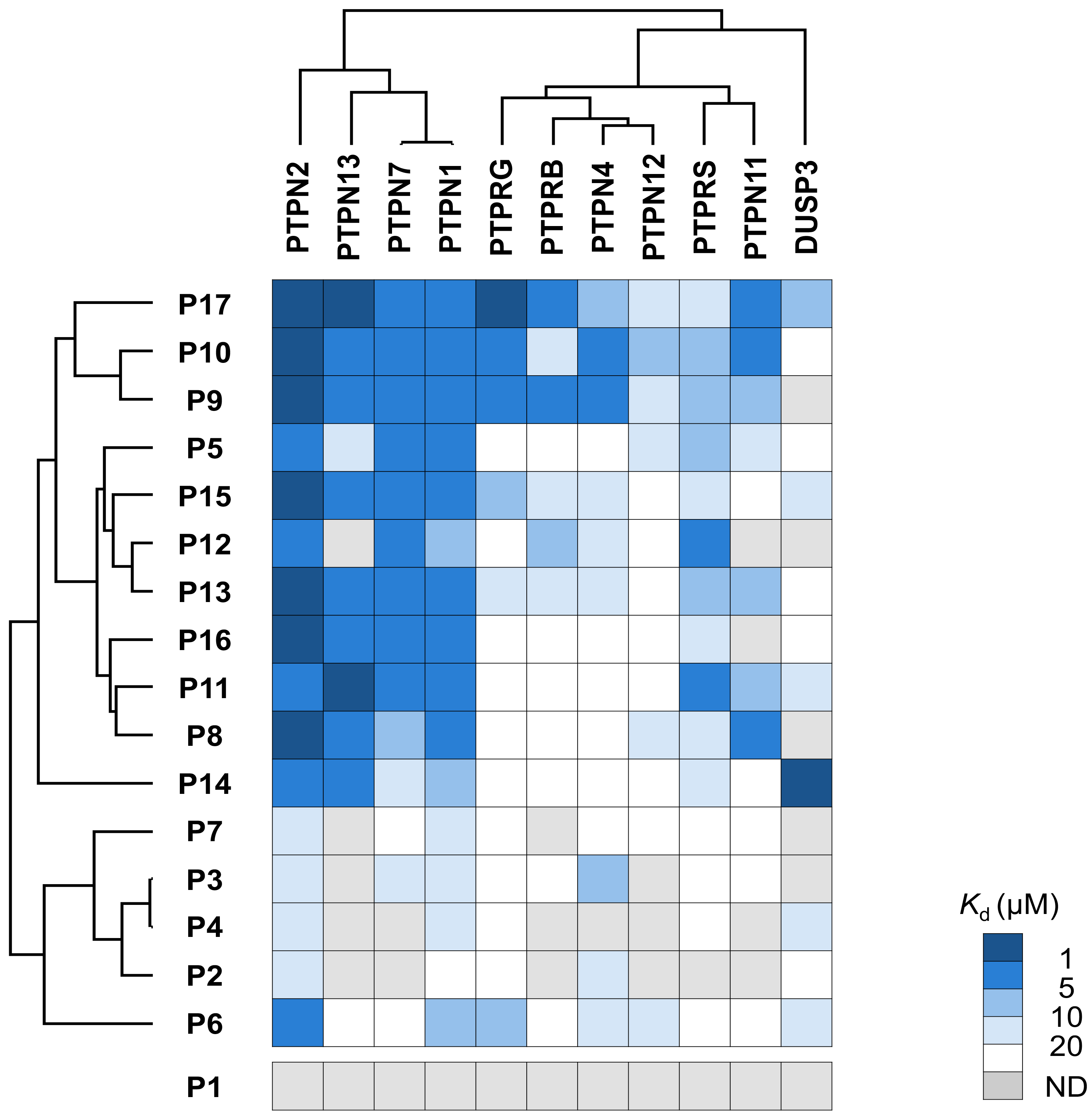
Binding affinity between F_2_Pmp probe and PTP. The dissociation constants between 17 F_2_Pmp probes and 11 PTPs are shown in the heat map. The PTPs and F_2_Pmp probes were clustered based on their dissociation constants.

Many PTPs showed very different *K_d_* values depending on the probe sequence. For example, many PTPs showed binding strength for **P9**, **P10**, and **P17** with *K_d_* values below 10 μM, while they did not show strong binding to **P4** and **P3** (Figure 1). Similar to the pulldown using cell extracts in our previous study^30^, most of the PTPs bound more strongly to **P17** than to **P13**∼**P16**, which were single amino acid substitutions of **P17**. Furthermore, none of the PTPs bound to F_2_Pmp alone (**P1**), and showed weak binding to **P2**.

We evaluated the binding selectivity of each F_2_Pmp probe to PTPs. The results showed that most of the F_2_Pmp probes were selectively bound to PTPN1, PTPN2, PTPN7, and PTPN13 (Figure S1C). On the other hand, the selectivity for other PTPs was different among the F_2_Pmp probes. For example, **P3** showed a preference for PTPN4. In addition, **P17** had strong selectivity for PTPRG, unlike its single amino acid substitutes, **P13**, **P14**, **P15**, and **P16**. Furthermore, **P14** showed strong selectivity for DUSP3 compared to the other four probes with single amino acid substitutions. These results indicate that the selectivity of the F_2_Pmp probe for PTPs is affected at the single amino acid level. In addition, in the cluster analysis, F_2_Pmp probes with one amino acid substitution were not classified close to each other, indicating that the difference of one amino acid has a significant effect on the selectivity for binding to PTPs.

DUSP3 showed a significant difference in the selectivity of probe sequence from classical PTP. This may indicate a difference in the binding selectivity of the F_2_Pmp probe for classical PTP and DUSP.

### A large-scale study on F_2_Pmp probe pulldown against cell lysates

We confirmed that the F_2_Pmp probe binds to many PTPs in a probe-sequence-dependent manner with a binding strength of several hundred nM to several μM *K_d_* value. Next, we evaluated whether endogenous PTPs are also enriched from cell extracts in a probe-sequence-dependent manner. F_2_Pmp peptide probes with biotin via a PEG_2_ linker at the N-terminus were bound to streptavidin resin and incubated with HEK293T or Jurkat cell extracts. After washing resin, proteins bound to the probes were eluted, digested, and subjected to LC/MS/MS analysis. To discriminate F_2_Pmp-dependent interacting proteins from non-specific interacting proteins, a peptide probe in which F_2_Pmp was replaced with tyrosine was used as a control probe.

From a total of 20 pulldown experiments using 17 probes and two cell extracts, a total of 5,507 proteins were quantified, of which 3,178 proteins were enriched as F_2_Pmp-dependent interacting proteins in any of the pulldown experiments (fold change > 2, p-value < 0.05) (Figure S2A). Of the 260 human phosphatases^31^, 93 were captured by any of the F_2_Pmp probes, 49 of which were enriched in an F_2_Pmp-dependent manner (Figure 2A). 14 of the 37 Classical PTPs encoded in the human genome were enriched (Figure S2B). Furthermore, as shown in Figure S2B, PTPN1 and PTPN2 were enriched in all F_2_Pmp probes except **P1**, but the enrichment profiles for other classical PTPs differed greatly from probe to probe. For example, **P9** and **P10** enriched various classical PTPs such as PTPN4 and PTPN11, while **P4**, **P7** and **P12** could enrich only PTPN1 and PTPN2. As we reported previously^30^, the enrichment profiles for PTPs in **P13** to **P17** differed greatly due to the effect of a single amino acid substitution. That is, five classical PTPs (PTPN1, PTPN2, PTPN13, PTPRF, and PTPRK) were enriched in **P13**, but only PTPN1 and PTPN2 were enriched in **P16**. Furthermore, in the comparison between **P13** and **P15**, while PTPN9, which could not be enriched in **P13**, could be enriched in **P15**, PTPRK could be enriched only in **P13**. The probe that could enrich the largest number of classical PTPs was **P17**, which could enrich 12 classical PTPs. This is consistent with the fact that **P17** showed strong binding to many recombinant PTPs (Figure S2B). **P2**, which is part of the probe sequence of **P13**, enriched only PTPN1 and PTPN2, while **P1** did not enrich any classical PTPs. These results indicate that the probe sequences contributed to the enrichment of PTP from cell extracts.

**Figure 2.**
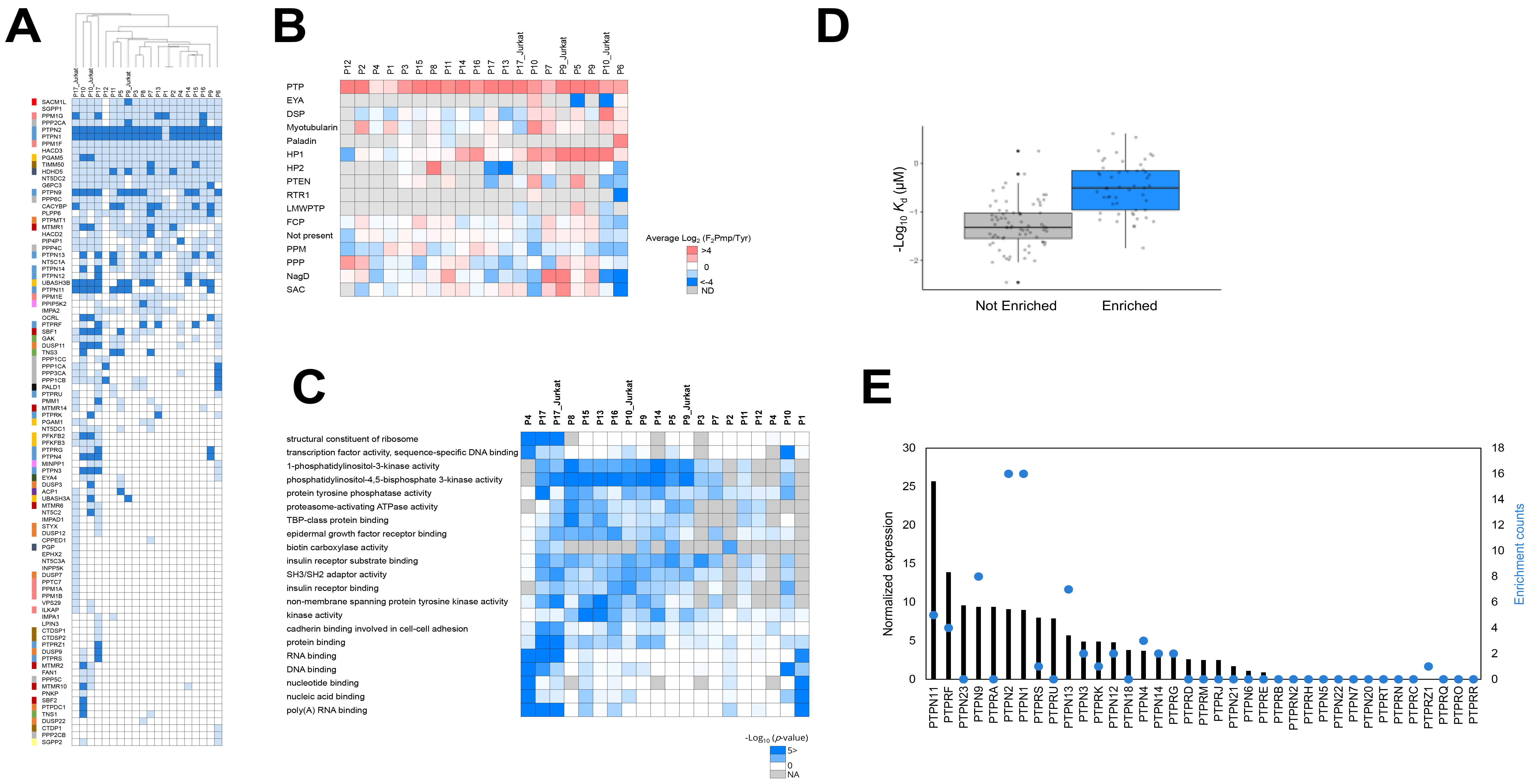
F_2_Pmp probe pulldown on cell extracts. A. Enrichment profile of phosphatase by F_2_Pmp probe. A heat map of phosphatases enriched from cell extracts by each F_2_Pmp probe is shown. Cells for enriched phosphatase are shown in dark blue, cells for unenriched but identified phosphatase are shown in light blue, and cells for unidentified phosphatase are shown in white. The pulldown conditions (probe and cell line) were clustered based on the enrichment profile. Phosphatases are also color-coded based on their family^33^. B. Enrichment profile of phosphatase family by F_2_Pmp probe. The enrichment ratio (F_2_Pmp probe/Y probe) for each phosphatase family is shown as a heat map on a log2 scale. The classification of phosphatase families is based on previous study^33^. C. GO enrichment analysis of proteins enriched by F_2_Pmp probe. The p-value (- log 10) of the GO term (Molecular Function) obtained by GO enrichment analysis for the proteins enriched by each F_2_Pmp probe is shown. GO enrichment analysis was performed by the Database for Annotation, Visualization and Integrated Discovery (DAVID)^63,64^. For the background dataset, all quantified proteins were used, and the minimum number of proteins was set to 2. Then, only GO terms that showed p value < 0.001 for two or more F_2_Pmp probes were extracted. D. Comparison between enrichment of PTP from HEK293T cell extracts by F_2_Pmp probe and binding affinity. The y-axis shows the dissociation constant (-log10) between each F_2_Pmp probe-PTP complex. The F_2_Pmp probe-PTP pair enriched from the HEK293T cell extract is designated as "Enriched" (blue), and the pair of F_2_Pmp probe and PTP not enriched is designated as "Not Enriched" (gray). E. Enrichment of PTP by F_2_Pmp probe and its expression level in HEK293 cells. The frequency with which each classical PTP was enriched from cell extracts by the F_2_Pmp probe is indicated by a blue dot. The expression level of each classical PTP in HEK293 cells (https://www.proteinatlas.org/) is shown as a bar graph.

In these 20 pulldown experiments, 16 PTPs other than classical PTPs, such as DUSP3, DUSP11 and UBASH3B^3^, were enriched by any of the F_2_Pmp probes. For phosphatases other than PTPs, 19 phosphatases such as PPP1CA, PPP3CA of the PPP family, TIMM50, HDHD5, and PGAM5 could be enriched by any of the F_2_Pmp probes. With regard to enriched phosphatases other than PTPs, it is known that TIMM50 exhibits tyrosine phosphatase activity in vitro^32^ and that the phosphatase domains of PGAM5 and HDHD5 are highly similar to those of PTPs, UBASH3A/B and EYA family respectively^33^. Therefore, these may have been enriched by the F_2_Pmp probe. Although the reason for the enrichment of other phosphatases other than PTPs requires further verification, these results indicate that the F_2_Pmp probe has the potential to enrich phosphatases other than PTPs. Next, we evaluated the differences in preference of F_2_Pmp probes among phosphatase families and found that classical PTP tended to be more strongly enriched in an F_2_Pmp-dependent manner than other phosphatase families (Figure 2B). One possible explanation for the higher enrichment of classical PTPs compared to other PTPs may be the difference of the substrate specificity for tyrosine or the size of the PTP domain^3^. As described above, the F_2_Pmp probe was able to enrich a variety of phosphatases, but tended to enrich more for classical PTPs.

In addition to phosphatases, SH2 domain-containing proteins were also enriched by the F_2_Pmp probe^34,35^. The SH2 domain is known to bind to the pY site in a sequence-dependent manner^36–39^. In fact, in our pulldown experiments, the SH2 domain-containing proteins were found to bind to the probe in a F_2_Pmp probe sequence-dependent manner (Figure S2C). We evaluated whether the sequence of the F_2_Pmp probe, which enriched SH2 domain, matched the known pY motif which is recognized by the SH2 domain^40^ and found that SH2 domain-containing proteins preferring the pY-X-X-P motif, such as CRKL and VAV2, were well enriched by the F_2_Pmp-X-X-P probe. On the other hand, SH2 domain-containing proteins preferring different pY motifs, such as PIK3R1 and YES, also showed specificity toward the F_2_Pmp-X-X- P probe. Further experiments are needed to conclude whether this is a result similar to the pulldown experiment using the pY probe or a result unique to F_2_Pmp. In addition, in a previous study^37^, SH2 domains showed the strong specificity for the amino acid sequence on the C-terminal side of the pY site. And in F_2_Pmp, **P8**∼**P17**, which have F_2_Pmp-(L/I)-(L/I)-P on the C-terminal side, showed a similar SH2 domain-containing protein enrichment profile, and cluster analysis showed that they were located near each other. On the other hand, the cluster analysis based on the enrichment profile of phosphatase (Figure 2A) did not show the same trend as SH2 domain-containing protein, indicating that the SH2 domain and phosphatase recognize different parts of the F_2_Pmp probe sequence.

As for the proteins enriched by the F_2_Pmp probe other than phosphatases and SH2 domain-containing proteins, GO enrichment analysis showed the proteins which were bound to nucleic acids such as RNA and DNA (Figure 2C). This is consistent with the results of pulldown experiments using pY peptides^39^, suggesting that proteins that bind to phosphate groups included in such as RNA and DNA may have been enriched by misrecognizing F_2_Pmp as a phosphate group. PKM and PRKCD have also been enriched by many F_2_Pmp probes, and it is known that PKM and PRKCD bind to pY peptides^41,42^, so they may have been similarly enriched by F_2_Pmp probes.

We then classified the F_2_Pmp peptide probe-PTP complexes based on their ability to be enriched from cell extracts, and evaluated the distribution of *K_d_* values among the F_2_Pmp peptide probe-PTP complexes for each class. As a result, the *K_d_* values between probes and PTP complexes enriched from cell extracts tended to be smaller than those between probes and PTP complexes that were not enriched (Figure 2D, S2D). Next, we evaluated the possibility that the PTP expression level in the cells could affect the enrichment by the F_2_Pmp probe. First, we evaluated the correlation between the mRNA expression level of classical PTP in HEK293 cells (https://www.proteinatlas.org/)43 and the number of F_2_Pmp probes that could enrich its classical PTP from HEK293T cell extracts (Figure 2E). The results showed that PTPs with very low expression levels (e.g. PTPN6, PTPRJ) were not enriched, suggesting that the frequency of enrichment was affected by the expression levels. In addition, the enrichment profiles of PTPs were different in the pulldown experiments using HEK293T cell extracts and Jurkat cell extracts (Figure S2B). For example, PTPN4, PTPN13 and PTPRG were enriched by **P9** from HEK293T cell extracts, but not from Jurkat cell extracts. According to the Human Protein Atlas, the expression levels of PTPRG and PTPN13 were lower in Jurkat cells than in HEK293 cells, suggesting the effect of expression level on enrichment. On the other hand, compared with PTPN2, PTPN9 and PTPN13, which were frequently enriched from HEK293T cell extracts, PTPN11 and PTPN12 were more abundantly expressed. One possibility is that PTPN2 and PTPN13 bound strongly to many F_2_Pmp probes as shown above, suggesting frequent enrichment. Taken together, our results indicate that classical PTPs are enriched from cell extracts depending on both the sequence of F_2_Pmp probes and the expression level of PTP. In our pulldown experiment with 17 F_2_Pmp probes, 13 of the 23 PTPs detected by RNA sequencing analysis in HEK293 cells were enriched (Figure S2E). In addition, PTPRZ1, which was not detected by RNA sequencing analysis, was also enriched. These results indicate that the F_2_Pmp probe can comprehensively enrich PTPs expressing in cells.

### PTP substrate specificity information

The pY substrate motif of PTP has been evaluated and confirmed by various methods^44–46^. Based on these findings, we hypothesized that the pY substrate motif information would correlate with PTP enrichment in pulldown experiments using the F_2_Pmp probe. Based on the in vitro substrate information of classical PTPs obtained by Palma et al^26^. We constructed PWMs and calculated PWM scores (Figure S3A)^47,48^, where PWM represents the probability of occurrence of amino acids in the vicinity of the pY site. Here, the columns of the matrix represent the relative positions from the pY site, and the rows represent the amino acids. The values of the matrix are log2 transformed probabilities of an amino acid appearing at a certain position. These probabilities are used to predict the putative substrates, and the PWM scores represent the certainty of those sequences as pY substrates.

First, we evaluated the relationship between PWM scores and *K_d_* values obtained from pulldown experiments using recombinant PTPs. The results showed that the PWM score was positively correlated with the inverse logarithm of the *K_d_* value (R=0.44) (Figure 3A). Next, we evaluated the relationship with respect to the pulldown experiment using cell extracts and found that the PWM score tended to be higher for the enriched PTP and F_2_Pmp probe pair (Figure 3B, S3B). These results confirm our hypothesis that the pY substrate motif information of PTP is correlated with the enrichment of PTP by the F_2_Pmp probe. Furthermore, these results support the validity of F_2_Pmp used as a mimetic of pY.

**Figure 3.**
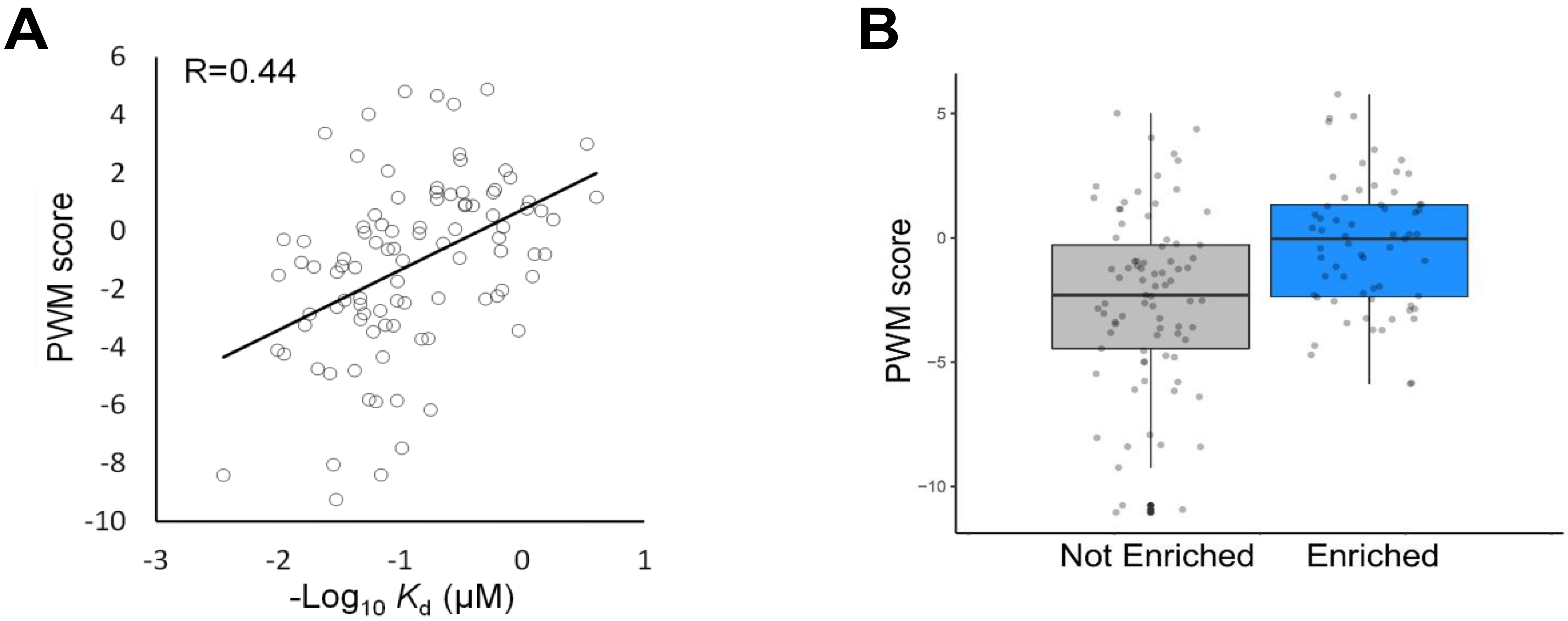
pY substrate specificity of PTP and PTP enrichment by F_2_Pmp probe. A. Comparison between binding affinity and PWM scores between classical PTP and F_2_Pmp Probes. The x-axis shows the dissociation constant (-log10) between each F_2_Pmp probe-PTP complex. The y-axis shows the PWM score between each F_2_Pmp probe and PTP. The correlation coefficients are calculated by the Pearson correlation coefficient. B. Comparison between enrichment of classical PTPs from HEK293T cell extracts by F_2_Pmp probe and PWM scores. The y-axis shows the PWM score between each F_2_Pmp probe and classical PTP. The pair of F_2_Pmp probe and classical PTP enriched from HEK293T cell extracts is designated as "Enriched" (blue), and the pair of F_2_Pmp probe and classical PTP not enriched is designated as "Not Enriched" (gray).

### PTP activity and binding to F_2_Pmp probe

The F_2_Pmp probe was then evaluated for its ability to identify PTPs competitively inhibited by PTP inhibitors. Orthovanadate binds reversibly to PTPs as a phosphate mimic and inhibits their activity^49^. We treated HEK293T cell extracts with orthovanadate (1.5 mM) and then performed pulldown experiments with **P9**. As a result, orthovanadate treatment significantly reduced the intensity of seven classical PTPs, including six classical PTPs that are known to bind to **P9** in an F_2_Pmp-dependent manner (Figure 4A). These results indicate that the F_2_Pmp probe discriminates the decrease in dephosphorylation activity caused by competitive inhibition of the active site of classical PTPs with endogenous substrates.

**Figure 4.**
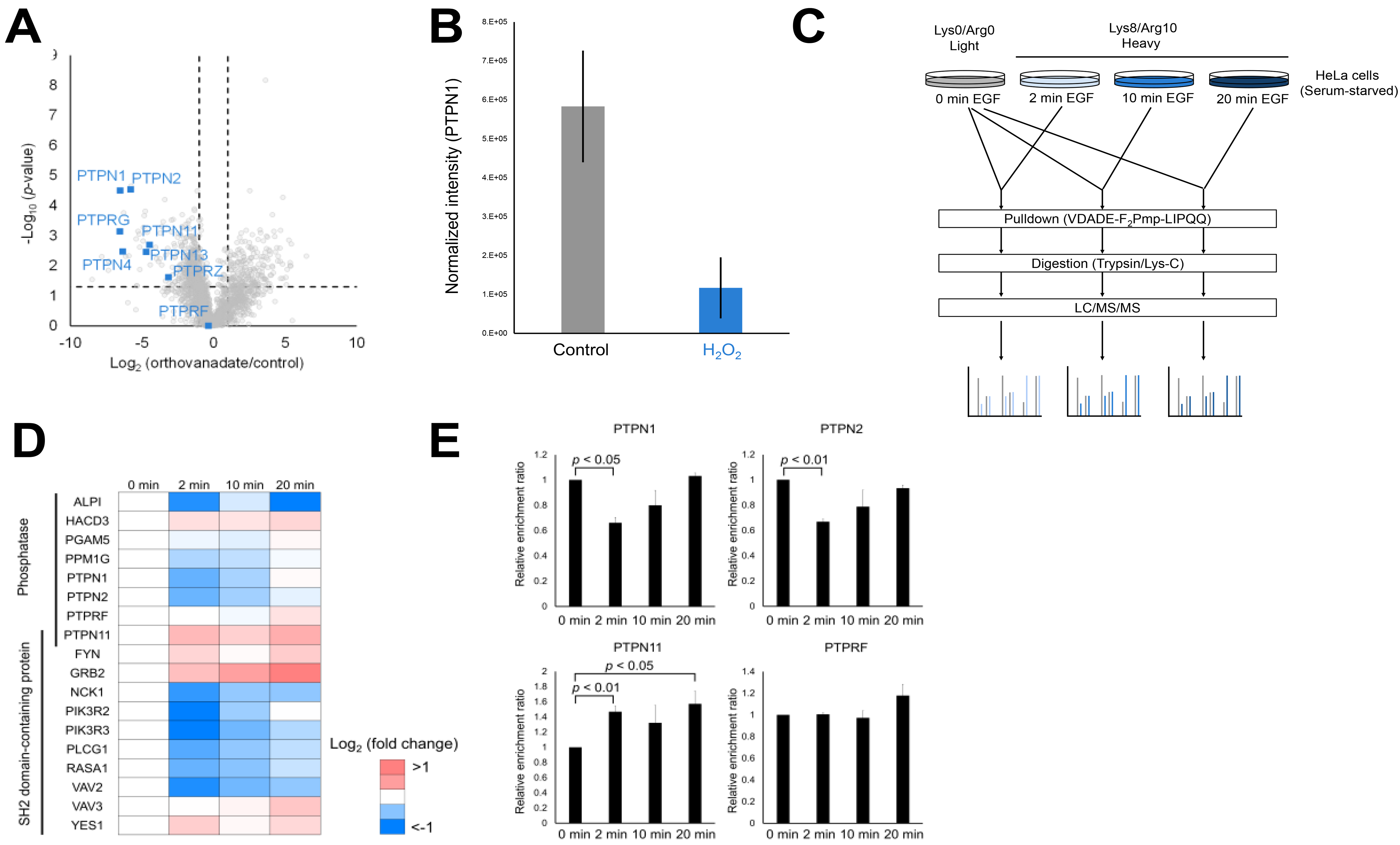
PTP activity and binding to F_2_Pmp probe. A. Competitive inhibition by orthovanadate and pulldown by F_2_Pmp probe. The x-axis indicates the enrichment ratio with and without orthovanadate addition. The p-value was calculated by Welch’s t-test. B. Oxidation by H_2_O_2_ and pulldown by F_2_Pmp probe. The normalized PTPN1 intensity with and without H_2_O_2_ treatment is shown. Pull downed PTPN1 intensity was normalized by the input PTPN1 intensity. C. Workflow of EGF stimulation and pulldown with F_2_Pmp probe D. The enrichment profiles of phosphatases and SH2 domain-containing proteins at each EGF stimulation time E. The enrichment ratio of PTP at each EGF stimulation time compared to the non-EGF treated sample. The p-value was calculated by one-sample t-test in Perseus^65^.

The cysteine residue in the active site of PTP is transiently oxidized by reactive oxygen species generated locally in the cell, resulting in an inactive form of PTP_50_. For example, it is well known that activation of EGFR causes a rapid increase in reactive oxygen species and inactivation of PTPN1^51,52^. Therefore, it is important to detect the inactivation of PTP by oxidation and Kumar et al. and Kalesh et al. have confirmed that the PTP probes developed by them did not react with PTP by H_2_O_2_ treatment in vitro^10,53^. Therefore, we evaluated whether the F_2_Pmp probe also discriminates the active state of PTP by oxidation. As in the previous study, we prepared PTPN1 treated with 1 mM H_2_O_2_ for 5 min and untreated PTPN1, incubated each with **P9**, and compared the enriched amount. In addition, a portion of the input was enzymatically digested and subjected to LC/MS/MS analysis. As a result, there was no difference in PTPN1 intensity between the H_2_O_2_-treated and control groups in the input, but in the pulldown sample, PTPN1 intensity was significantly decreased in the H_2_O_2_-treated group (Figure 4B). These results indicate that the F_2_Pmp probe discriminates PTP inactivated by oxidation.

We then used the F_2_Pmp probe to evaluate the activity variation of PTP in the cell. In this experiment, EGF was treated to stable isotope-labeled HeLa cells for 2, 10, or 20 minutes (Figure 4C). After mixing the extracts of the untreated cells and each EGF-treated cells, pulldown by **P9**, digested, and analyzed by LC/MS/MS. As a result, 448 proteins, including 8 phosphatases and 11 SH2 domain-containing proteins, were quantified. Out of them, 101 proteins, including 4 phosphatases and 8 SH2 domain-containing proteins, were changed significantly in dependence on EGF stimulation (H/L ratio > 0.5 or < −0.5, p-value < 0.05) (Figure S4). The enrichment of PTPN1 decreased 2 min after EGF treatment and then increased, which is consistent with the previous report (Figure 4D, E)^51^. Furthermore, PTPN2 behaved similarly to PTPN1, indicating that the activity of PTPN2 also fluctuated in an EGF stimulation-dependent manner. PTPN11 also showed EGF-stimulated dependent fluctuations in enrichment. However, since PTPN11 is composed of a PTP domain and two SH2 domains, it is difficult to estimate the binding domain of this probe to PTPN11, and further verification is needed to determine whether the variation in enrichment of PTPN11 reflects PTP activity. For PTPRF, there was no significant change in enrichment, indicating no change of the activity upon EGF stimulation. In addition to PTPs, SH2 domain-related proteins, such as RASA1 and PI3K complex, were transiently decreased in enrichment after EGF stimulation (Figure 4D). In EGF stimulation, a transient increase in the amount of pY sites in the cell is caused by the activation of tyrosine kinases^38,54^. SH2 domains bind to these pY sites and form a complex that transmits signals downstream^38^. Therefore, one possibility is that EGF stimulation changes the amount of SH2 domains that can be bound by the F_2_Pmp probe in cell extracts, and thus the enrichment of SH2 domain-containing proteins. We then perform pathway enrichment analysis to characterize the proteins that were fluctuated in this pull-down. Pathway enrichment analysis showed that the proteins fluctuated in EGF stimulation were mapped to the pathways well known to be associated with EGF stimulation, such as ROS or ErbB signaling pathway (Figure S4). Proteins on these pathways are known to show various responses to EGF stimulation. For example, ATP synthase (ATP5F1A, ATP5F1B, ATP5F1C, ATP5MF), which was altered in this pull-down, is attributed to the ROS pathway and it is known that ATP turnover and ATP citrate lyase activity increase upon EGF stimulation^55,56^. And, proteins such as G6PD, PFKP and PFKM, which were attributed to the central carbon metabolism in cancer, are tightly linked to glycolysis, which is stimulated by EGF. In these cases, further experiments are needed to determine what the enrichment of ATP synthase and glycolysis-related proteins in the F_2_Pmp probe pull-down represents in biological meanings, but the enrichment may be altered by changes of the protein (structure, activity, complex) caused by EGF stimulation.

## CONCLUSION

We aimed to develop a comprehensive enrichment method for PTP and evaluated the relationship between probe sequence and enriched PTP, and the correlation between enrichment by F_2_Pmp probe and PTP activity. In this study, we showed that the F_2_Pmp probe binds to PTP in a sequence-dependent and PTP activity-dependent manner. In addition, this study is the first successful example of comprehensive enrichment of classical PTP at the protein level.

By systematically designing the F_2_Pmp probe sequence and increasing the sequence diversity, we have shown that not only could PTPs be comprehensively enriched, but also some serine threonine phosphatases could be enriched. These results raise the hope that comprehensive enrichment of human phosphatases will be possible by optimizing the probe sequence and using non-hydrolyzable analogs of pS and pT.

Although F_2_Pmp is used as a mimetic of pY, it differs from pY in size and bond length^57^. Therefore, it was unclear whether the pY substrate specificity information of PTP could be used for F_2_Pmp probes. In this study, we obtained large-scale information on PTP enrichment by F_2_Pmp probes, which enabled us to conduct a detailed analysis of the correlation between PTP enrichment by F_2_Pmp probes and pY substrate specificity information of PTP. We believe that this study has enabled the efficient design of F_2_Pmp probe sequences.

In addition, our approach was capable of PTP enrichment at the protein level and detection of competitive inhibition by orthovanadate. Therefore, this method could be applied to obtain the binding profile of inhibitors to PTPs, as performed for kinase inhibitors using kinobeads.

It is well known that the activity of tyrosine kinases changes in response to external stimuli and regulates downstream phosphorylation signals^58^. On the other hand, there are only a limited number of reports on the regulation of PTP activity^51,59–62^. In this study, we showed that the activity of not only the known PTPN1 but also PTPN2 fluctuates upon EGF stimulation. This result further supports that intracellular phosphorylation signals are tightly regulated not only by kinases but also by phosphatases. The F_2_Pmp probe is expected to be a useful probe that can comprehensively capture the changes in the activity of these PTPs.

## Author Contributions

T.N. and Y.I. designed research. T.N. performed research and analyzed data. T.N. and Y.I. wrote the paper.

---

**Table S1.**
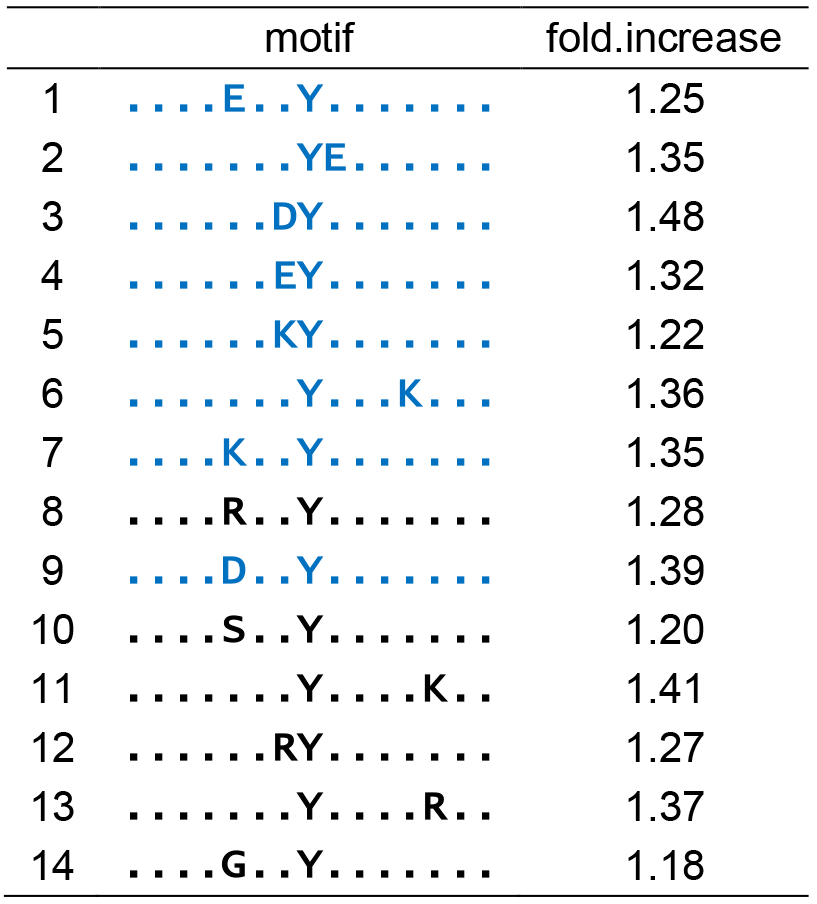
Enriched motifs around the pY site registered in PhosphoSitePlus. The motifs contained in the probes used in this study are indicated in blue.

**Figure S1.**
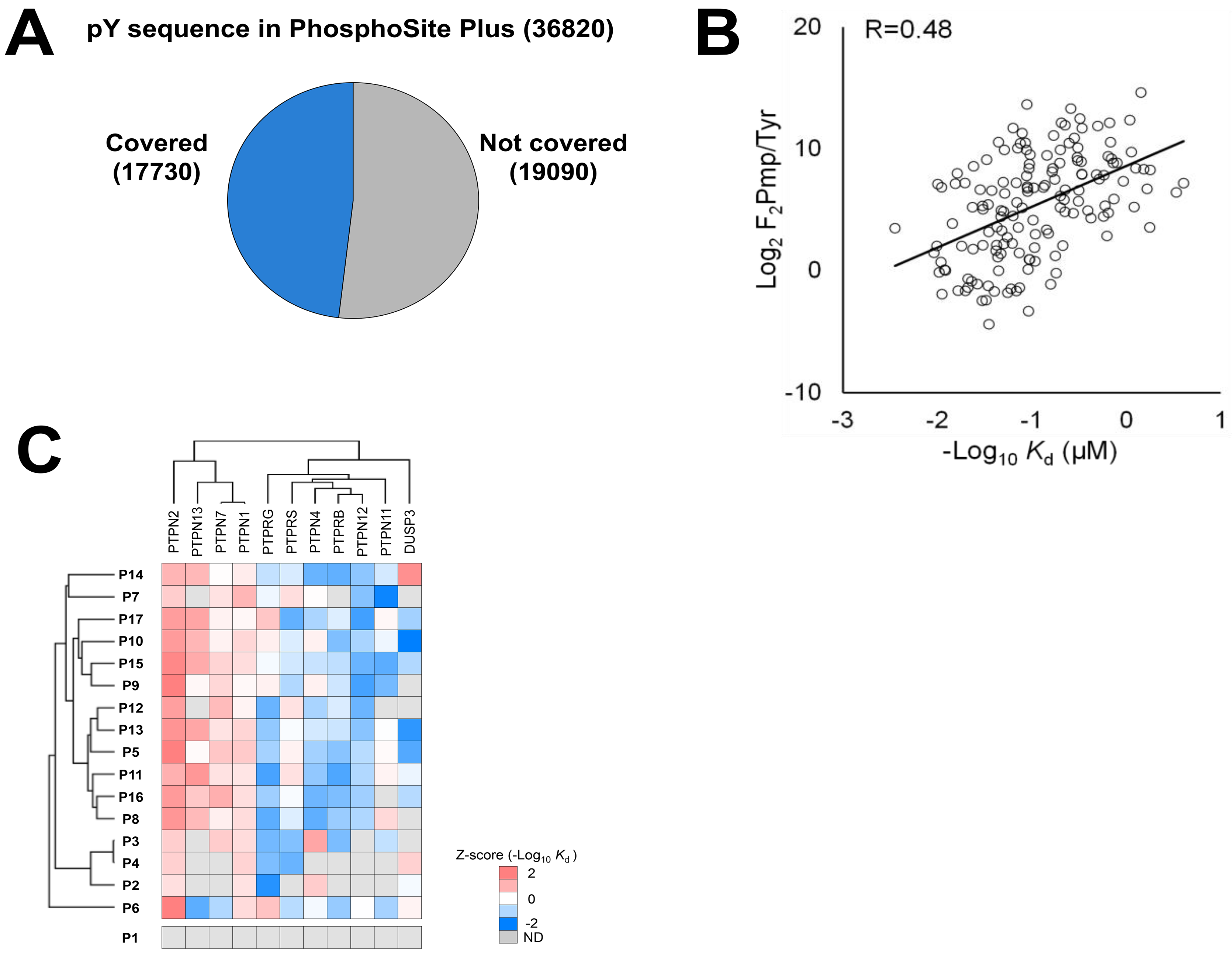
Binding affinity between F_2_Pmp probe and PTP. A. Percentage of pY sites with motifs shown in blue in Table S1 B. Comparison of binding affinity and enrichment ratio for the F_2_Pmp probes-PTP complex. The x-axis shows the dissociation constant (-log10) between each F_2_Pmp probe-PTP complex. The y-axis shows the enrichment ratio of proteins in pulldown experiments with each probe (F_2_Pmp probe/Y probe) on a log2 scale. The correlation coefficients were calculated by Pearson correlation coefficient. C. Binding selectivity between F_2_Pmp probe and PTP. The dissociation constant (log 10) at Figure 1A is normalized for each F_2_Pmp probe (binding selectivity). The heat map shows the z-score (binding selectivity) for each F_2_Pmp probe. The PTP and F_2_Pmp probes were clustered based on their binding selectivity.

**Figure S2.**
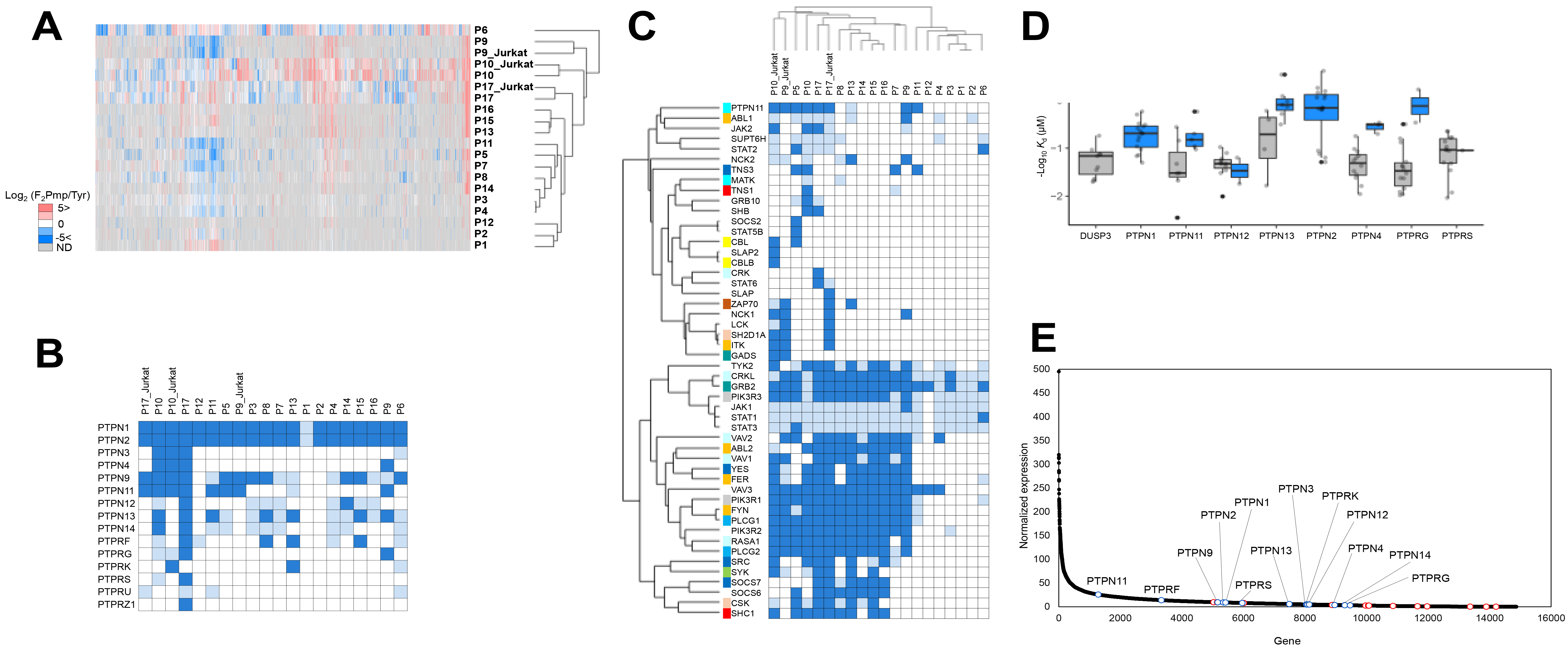
F_2_Pmp probe pulldown on cell extracts. A. F_2_Pmp probe pulldown on cell extracts. The enrichment ratio (F_2_Pmp probe/Y probe) for cell extracts by each F_2_Pmp probe is shown on a log2 scale. The pulldown conditions (probe and cell type) were clustered based on the enrichment ratio. B. Enrichment profile of classical PTPs by F_2_Pmp probe. A heat map of classical PTPs enriched from cell extracts by each F_2_Pmp probe is shown. Cells for enriched classical PTPs are shown in dark blue, cells for unenriched but identified classical PTPs are shown in light blue, and cells for unidentified classical PTPs are shown in white. C. Enrichment profile of SH2 domain-containing proteins by F_2_Pmp probe. SH2 domain-containing proteins enriched from cell extracts by each F_2_Pmp probe are shown in the heat map. Cells for enriched SH2 domain-containing proteins are shown in dark blue, cells for SH2 domain-containing proteins that were not enriched but were identified are shown in light blue, and cells for SH2 domain-containing proteins that were not identified are shown in white. Pulldown conditions (probe and cell type) and SH2 domain-containing proteins were clustered based on enrichment profiles. SH2 domain-containing proteins are color-coded based on their sequence specificity to the pY peptide^37^. D. Comparison between enrichment of PTP from HEK293T cell extracts by F_2_Pmp probe and binding affinity. The y-axis shows the dissociation constant (-log10) between each F_2_Pmp probe-PTP complex. The F_2_Pmp probe-PTP pair enriched from the HEK293T cell extract is designated as "Enriched" (blue), and the pair of F_2_Pmp probe and PTP not enriched is designated as "Not Enriched" (gray). They are represented for each PTP. E. The expression levels in HEK293 cells (https://www.proteinatlas.org/) are shown as black dots. Classical PTPs enriched from cell extracts by the F_2_Pmp probe are shown as blue dots, and classical PTPs not enriched are shown as red dots.

**Figure S3.**
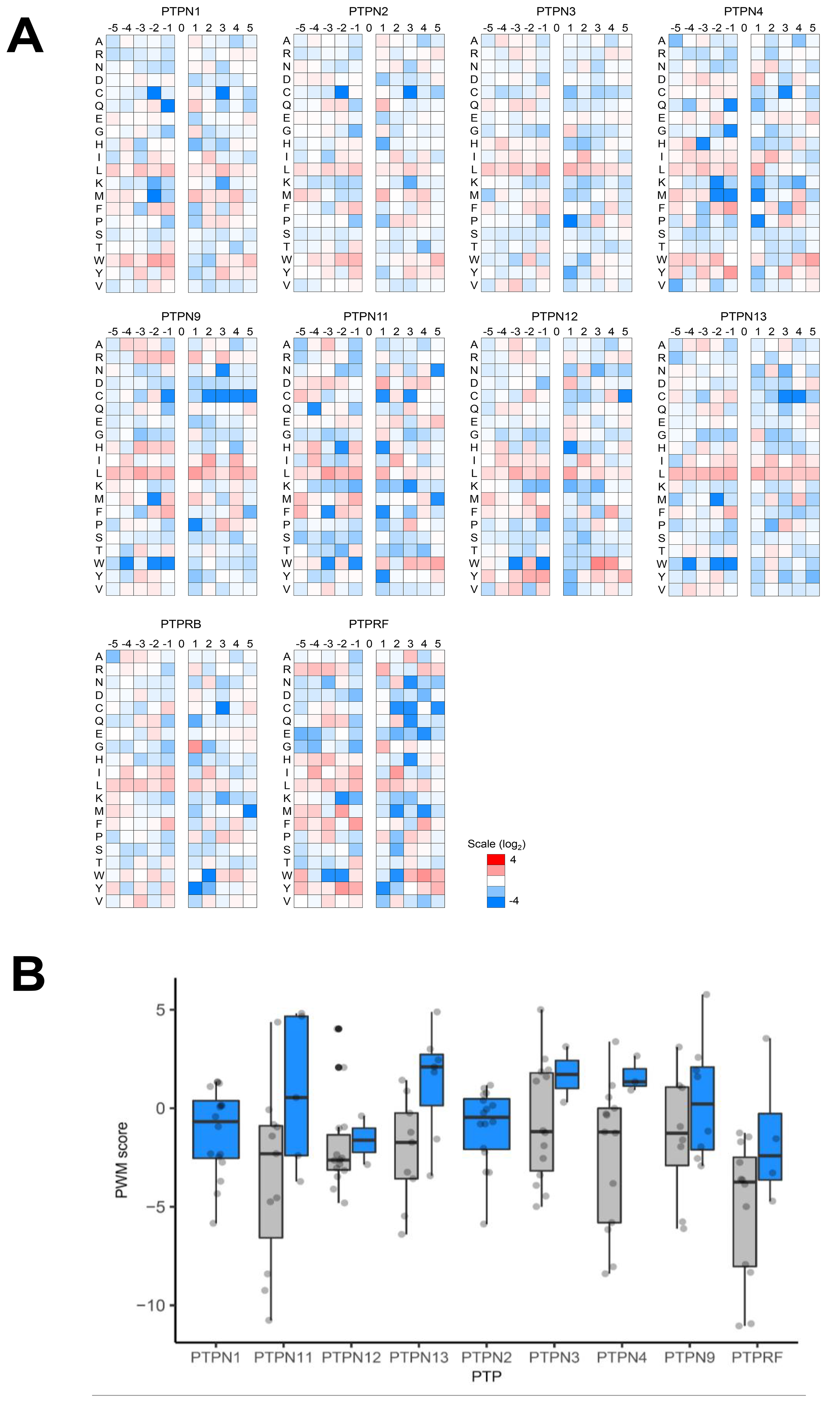
pY substrate specificity of PTP and PTP enrichment by F_2_Pmp probe. A. PWMs of each classical PTP B. Comparison between enrichment of classical PTPs from HEK293T cell extracts by F_2_Pmp probe and PWM scores. The y-axis shows the PWM score between each F_2_Pmp probe and classical PTP. The pair of F_2_Pmp probe and classical PTP enriched from HEK293T cell extracts is designated as "Enriched" (blue), and the pair of F_2_Pmp probe and classical PTP not enriched is designated as "Not Enriched" (gray). They are represented for each classical PTP.

**Figure S4.**
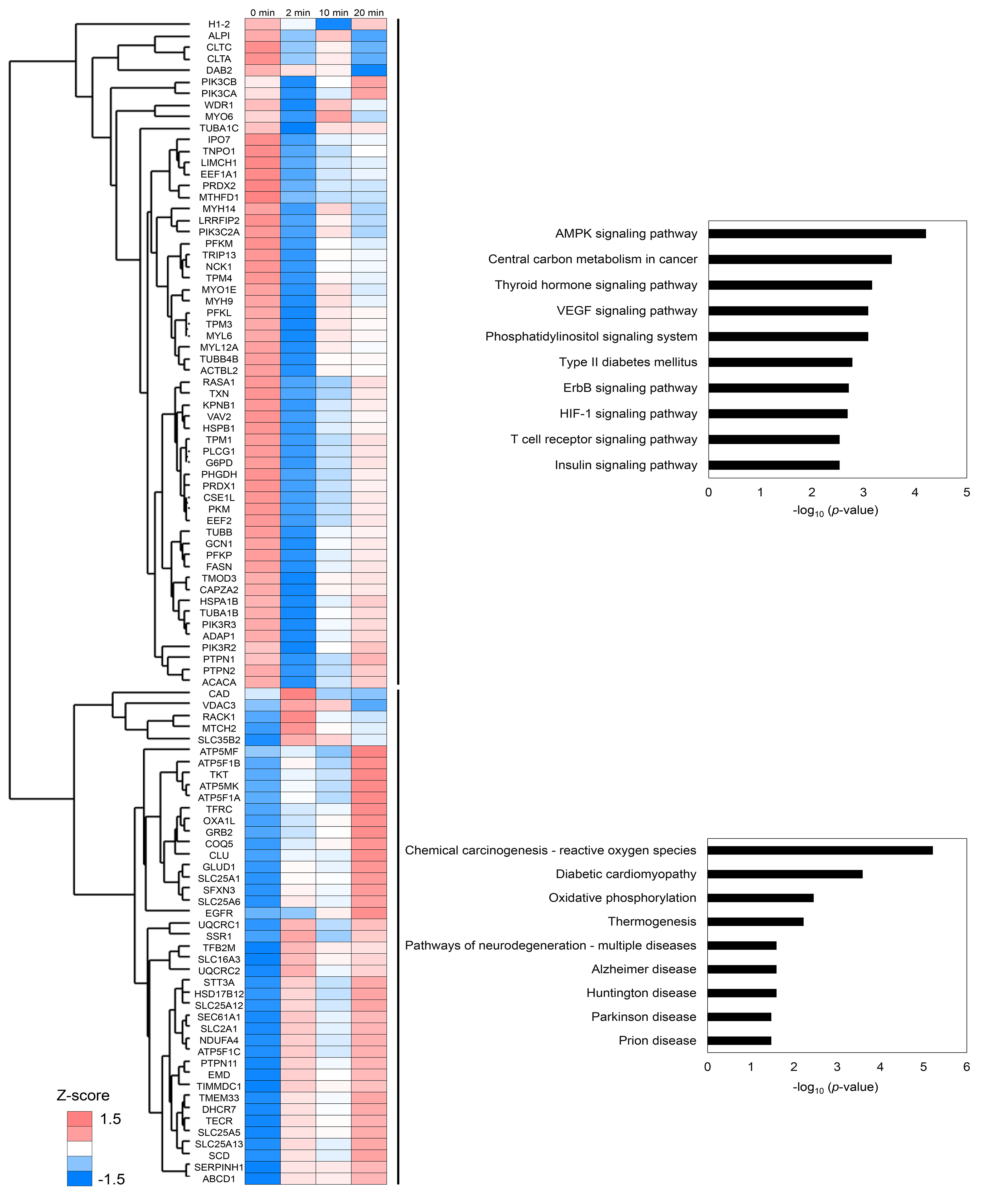
EGF stimulation and pull-down with F_2_Pmp probe. The enrichment profiles of proteins significantly fluctuating at each EGF stimulation time. Bar graph represents the p-value of the significantly enriched pathway (p-value <0.05, TOP10) in DAVID.

## References

(1) Julien, S. G.; Dubé, N.; Hardy, S.; Tremblay, M. L. Inside the Human Cancer Tyrosine Phosphatome. Nat. Rev. Cancer 2011, 11 (1), 35–49.

(2) Saraon, P.; Pathmanathan, S.; Snider, J.; Lyakisheva, A.; Wong, V.; Stagljar, I. Receptor Tyrosine Kinases and Cancer: Oncogenic Mechanisms and Therapeutic Approaches. Oncogene 2021, 40 (24), 4079–4093.

(3) Alonso, A.; Pulido, R. The Extended Human PTPome: A Growing Tyrosine Phosphatase Family. FEBS J. 2016, 283 (11), 2197–2201.

(4) Bantscheff, M.; Eberhard, D.; Abraham, Y.; Bastuck, S.; Boesche, M.; Hobson, S.; Mathieson, T.; Perrin, J.; Raida, M.; Rau, C.; Reader, V.; Sweetman, G.; Bauer, A.; Bouwmeester, T.; Hopf, C.; Kruse, U.; Neubauer, G.; Ramsden, N.; Rick, J.; Kuster, B.; Drewes, G. Quantitative Chemical Proteomics Reveals Mechanisms of Action of Clinical ABL Kinase Inhibitors. Nat. Biotechnol. 2007, 25 (9), 1035–1044.

(5) Lyons, S. P.; Jenkins, N. P.; Nasa, I.; Choy, M. S.; Adamo, M. E.; Page, R.; Peti, W.; Moorhead, G. B.; Kettenbach, A. N. A Quantitative Chemical Proteomic Strategy for Profiling Phosphoprotein Phosphatases from Yeast to Humans. Mol. Cell. Proteomics 2018, 17 (12), 2448–2461.

(6) Bantscheff, M.; Eberhard, D.; Abraham, Y.; Bastuck, S.; Boesche, M.; Hobson, S.; Mathieson, T.; Perrin, J.; Raida, M.; Rau, C.; Reader, V.; Sweetman, G.; Bauer, A.; Bouwmeester, T.; Hopf, C.; Kruse, U.; Neubauer, G.; Ramsden, N.; Rick, J.; Kuster, B.; Drewes, G. Quantitative Chemical Proteomics Reveals Mechanisms of Action of Clinical ABL Kinase Inhibitors. Nat. Biotechnol. 2007, 25 (9), 1035–1044.

(7) Klaeger, S.; Heinzlmeir, S.; Wilhelm, M.; Polzer, H.; Vick, B.; Koenig, P.-A.; Reinecke, M.; Ruprecht, B.; Petzoldt, S.; Meng, C.; Zecha, J.; Reiter, K.; Qiao, H.; Helm, D.; Koch, H.; Schoof, M.; Canevari, G.; Casale, E.; Depaolini, S. R.; Feuchtinger, A.; Wu, Z.; Schmidt, T.; Rueckert, L.; Becker, W.; Huenges, J.; Garz, A.-K.; Gohlke, B.-O.; Zolg, D. P.; Kayser, G.; Vooder, T.; Preissner, R.; Hahne, H.; Tõnisson, N.; Kramer, K.; Götze, K.; Bassermann, F.; Schlegl, J.; Ehrlich, H.-C.; Aiche, S.; Walch, A.; Greif, P. A.; Schneider, S.; Felder, E. R.; Ruland, J.; Médard, G.; Jeremias, I.; Spiekermann, K.; Kuster, B. The Target Landscape of Clinical Kinase Drugs. Science 2017, 358 (6367). https://doi.org/10.1126/science.aan4368.

(8) Lyons, S. P.; Jenkins, N. P.; Nasa, I.; Choy, M. S.; Adamo, M. E.; Page, R.; Peti, W.; Moorhead, G. B.; Kettenbach, A. N. A Quantitative Chemical Proteomic Strategy for Profiling Phosphoprotein Phosphatases from Yeast to Humans. Mol. Cell. Proteomics 2018, 17 (12), 2448–2461.

(9) Karisch, R.; Fernandez, M.; Taylor, P.; Virtanen, C.; St-Germain, J. R.; Jin, L. L.; Harris, I. S.; Mori, J.; Mak, T. W.; Senis, Y. A.; Östman, A.; Moran, M. F.; Neel, B. G. Global Proteomic Assessment of the Classical Protein-Tyrosine Phosphatome and “Redoxome.” Cell 2011, 146 (5), 826–840.

(10) Kumar, S.; Zhou, B.; Liang, F.; Yang, H.; Wang, W.-Q.; Zhang, Z.-Y. Global Analysis of Protein Tyrosine Phosphatase Activity with Ultra-Sensitive Fluorescent Probes. J. Proteome Res. 2006, 5 (8), 1898–1905.

(11) Liu, S.; Zhou, B.; Yang, H.; He, Y.; Jiang, Z.-X.; Kumar, S.; Wu, L.; Zhang, Z.-Y. Aryl Vinyl Sulfonates and Sulfones as Active Site-Directed and Mechanism-Based Probes for Protein Tyrosine Phosphatases. J. Am. Chem. Soc. 2008, 130 (26), 8251–8260.

(12) Kumar, S.; Zhou, B.; Liang, F.; Wang, W.-Q.; Huang, Z.; Zhang, Z.-Y. Activity-Based Probes for Protein Tyrosine Phosphatases. Proc. Natl. Acad. Sci. U. S. A. 2004, 101 (21), 7943–7948.

(13) Lo, L.-C.; Pang, T.-L.; Kuo, C.-H.; Chiang, Y.-L.; Wang, H.-Y.; Lin, J.-J. Design and Synthesis of Class-Selective Activity Probes for Protein Tyrosine Phosphatases. J. Proteome Res. 2002, 1 (1), 35–40.

(14) Kalesh, K. A.; Tan, L. P.; Lu, K.; Gao, L.; Wang, J.; Yao, S. Q. Peptide-Based Activity-Based Probes (ABPs) for Target-Specific Profiling of Protein Tyrosine Phosphatases (PTPs). Chem. Commun. 2010, 46 (4), 589–591.

(15) Ge, J.; Cheng, X.; Tan, L. P.; Yao, S. Q. Ugi Reaction-Assisted Rapid Assembly of Affinity-Based Probes against Potential Protein Tyrosine Phosphatases. Chem. Commun. 2012, 48 (37), 4453–4455.

(16) Casey, G. R.; Stains, C. I. Interrogating Protein Phosphatases with Chemical Activity Probes. Chemistry 2018, 24 (31), 7810–7824.

(17) Chen, L.; Wu, L.; Otaka, A.; Smyth, M. S.; Roller, P. P.; Burke, T. R. Jr; den Hertog, J.; Zhang, Z. Y. Why Is Phosphonodifluoromethyl Phenylalanine a More Potent Inhibitory Moiety than Phosphonomethyl Phenylalanine toward Protein-Tyrosine Phosphatases? Biochem. Biophys. Res. Commun. 1995, 216 (3), 976–984.

(18) Meyer, C.; Hoeger, B.; Chatterjee, J.; Köhn, M. Azide-Alkyne Cycloaddition-Mediated Cyclization of Phosphonopeptides and Their Evaluation as PTP1B Binders and Enrichment Tools. Bioorg. Med. Chem. 2015, 23 (12), 2848–2853.

(19) Meyer, C.; Hoeger, B.; Temmerman, K.; Tatarek-Nossol, M.; Pogenberg, V.; Bernhagen, J.; Wilmanns, M.; Kapurniotu, A.; Köhn, M. Development of Accessible Peptidic Tool Compounds To Study the Phosphatase PTP1B in Intact Cells. ACS Chemical Biology. 2014, pp 769–776. https://doi.org/10.1021/cb400903u.

(20) Tsumagari, K.; Niinae, T.; Otaka, A.; Ishihama, Y. Peptide Probes Containing a Non-Hydrolyzable Phosphotyrosine-Mimetic Residue for Enrichment of Protein Tyrosine Phosphatases. Proteomics 2021, e2100144.

(21) Masuda, T.; Tomita, M.; Ishihama, Y. Phase Transfer Surfactant-Aided Trypsin Digestion for Membrane Proteome Analysis. J. Proteome Res. 2008, 7 (2), 731–740.

(22) Rappsilber, J.; Mann, M.; Ishihama, Y. Protocol for Micro-Purification, Enrichment, Pre-Fractionation and Storage of Peptides for Proteomics Using StageTips. Nat. Protoc. 2007, 2 (8), 1896–1906.

(23) Cox, J.; Mann, M. MaxQuant Enables High Peptide Identification Rates, Individualized P.p.b.-Range Mass Accuracies and Proteome-Wide Protein Quantification. Nature Biotechnology. 2008, pp 1367–1372. https://doi.org/10.1038/nbt.1511.

(24) Tyanova, S.; Temu, T.; Sinitcyn, P.; Carlson, A.; Hein, M. Y.; Geiger, T.; Mann, M.; Cox, J. The Perseus Computational Platform for Comprehensive Analysis of (prote)omics Data. Nat. Methods 2016, 13 (9), 731–740.

(25) Sharma, K.; Weber, C.; Bairlein, M.; Greff, Z.; Kéri, G.; Cox, J.; Olsen, J. V.; Daub, H. Proteomics Strategy for Quantitative Protein Interaction Profiling in Cell Extracts. Nat. Methods 2009, 6 (10), 741–744.

(26) Palma, A.; Tinti, M.; Paoluzi, S.; Santonico, E.; Brandt, B. W.; Hooft van Huijsduijnen, R.; Masch, A.; Heringa, J.; Schutkowski, M.; Castagnoli, L.; Cesareni, G. Both Intrinsic Substrate Preference and Network Context Contribute to Substrate Selection of Classical Tyrosine Phosphatases. J. Biol. Chem. 2017, 292 (12), 4942–4952.

(27) Wagih, O.; Sugiyama, N.; Ishihama, Y.; Beltrao, P. Uncovering Phosphorylation-Based Specificities through Functional Interaction Networks. Mol. Cell. Proteomics 2016, 15 (1), 236–245.

(28) Hornbeck, P. V.; Zhang, B.; Murray, B.; Kornhauser, J. M.; Latham, V.; Skrzypek, E. PhosphoSitePlus, 2014: Mutations, PTMs and Recalibrations. Nucleic Acids Res. 2015, 43 (Database issue), D512–D520.

(29) Zhang, Y. L.; Yao, Z. J.; Sarmiento, M.; Wu, L.; Burke, T. R., Jr; Zhang, Z. Y. Thermodynamic Study of Ligand Binding to Protein-Tyrosine Phosphatase 1B and Its Substrate-Trapping Mutants. J. Biol. Chem. 2000, 275 (44), 34205–34212.

(30) Tsumagari, K.; Niinae, T.; Otaka, A.; Ishihama, Y. Peptide Probes Containing a Non-Hydrolyzable Phosphotyrosine-Mimetic Residue for Enrichment of Protein Tyrosine Phosphatases. Proteomics 2021, e2100144.

(31) Damle, N. P.; Köhn, M. The Human DEPhOsphorylation Database DEPOD: 2019 Update. Database 2019, 2019. https://doi.org/10.1093/database/baz133.

(32) Guo, Y.; Cheong, N.; Zhang, Z.; De Rose, R.; Deng, Y.; Farber, S. A.; Fernandes-Alnemri, T.; Alnemri, E. S. Tim50, a Component of the Mitochondrial Translocator, Regulates Mitochondrial Integrity and Cell Death. J. Biol. Chem. 2004, 279 (23), 24813–24825.

(33) Chen, M. J.; Dixon, J. E.; Manning, G. Genomics and Evolution of Protein Phosphatases. Sci. Signal. 2017, 10 (474). https://doi.org/10.1126/scisignal.aag1796.

(34) Tsumagari, K.; Niinae, T.; Otaka, A.; Ishihama, Y. Peptide Probes Containing a Non-Hydrolyzable Phosphotyrosine-Mimetic Residue for Enrichment of Protein Tyrosine Phosphatases. Proteomics 2021, e2100144.

(35) Burke, T. R., Jr; Smyth, M. S.; Otaka, A.; Nomizu, M.; Roller, P. P.; Wolf, G.; Case, R.; Shoelson, S. E. Nonhydrolyzable Phosphotyrosyl Mimetics for the Preparation of Phosphatase-Resistant SH2 Domain Inhibitors. Biochemistry 1994, 33 (21), 6490–6494.

(36) Schulze, W. X.; Deng, L.; Mann, M. Phosphotyrosine Interactome of the ErbB-Receptor Kinase Family. Mol. Syst. Biol. 2005, 1, 2005.0008.

(37) Tinti, M.; Kiemer, L.; Costa, S.; Miller, M. L.; Sacco, F.; Olsen, J. V.; Carducci, M.; Paoluzi, S.; Langone, F.; Workman, C. T.; Blom, N.; Machida, K.; Thompson, C. M.; Schutkowski, M.; Brunak, S.; Mann, M.; Mayer, B. J.; Castagnoli, L.; Cesareni, G. The SH2 Domain Interaction Landscape. Cell Rep. 2013, 3 (4), 1293–1305.

(38) Lundby, A.; Franciosa, G.; Emdal, K. B.; Refsgaard, J. C.; Gnosa, S. P.; Bekker-Jensen, D. B.; Secher, A.; Maurya, S. R.; Paul, I.; Mendez, B. L.; Kelstrup, C. D.; Francavilla, C.; Kveiborg, M.; Montoya, G.; Jensen, L. J.; Olsen, J. V. Oncogenic Mutations Rewire Signaling Pathways by Switching Protein Recruitment to Phosphotyrosine Sites. Cell 2019, 179 (2), 543–560.e26.

(39) Hanke, S.; Mann, M. The Phosphotyrosine Interactome of the Insulin Receptor Family and Its Substrates IRS-1 and IRS-2. Mol. Cell. Proteomics 2009, 8 (3), 519– 534.

(40) Tinti, M.; Kiemer, L.; Costa, S.; Miller, M. L.; Sacco, F.; Olsen, J. V.; Carducci, M.; Paoluzi, S.; Langone, F.; Workman, C. T.; Blom, N.; Machida, K.; Thompson, C. M.; Schutkowski, M.; Brunak, S.; Mann, M.; Mayer, B. J.; Castagnoli, L.; Cesareni, G. The SH2 Domain Interaction Landscape. Cell Rep. 2013, 3 (4), 1293–1305.

(41) Benes, C. H.; Wu, N.; Elia, A. E. H.; Dharia, T.; Cantley, L. C.; Soltoff, S. P. The C2 Domain of PKCdelta Is a Phosphotyrosine Binding Domain. Cell 2005, 121 (2), 271–280.

(42) Christofk, H. R.; Vander Heiden, M. G.; Wu, N.; Asara, J. M.; Cantley, L. C. Pyruvate Kinase M2 Is a Phosphotyrosine-Binding Protein. Nature 2008, 452 (7184), 181–186.

(43) Uhlén, M.; Fagerberg, L.; Hallström, B. M.; Lindskog, C.; Oksvold, P.; Mardinoglu, A.; Sivertsson, Å.; Kampf, C.; Sjöstedt, E.; Asplund, A.; Olsson, I.; Edlund, K.; Lundberg, E.; Navani, S.; Szigyarto, C. A.-K.; Odeberg, J.; Djureinovic, D.; Takanen, J. O.; Hober, S.; Alm, T.; Edqvist, P.-H.; Berling, H.; Tegel, H.; Mulder, J.; Rockberg, J.; Nilsson, P.; Schwenk, J. M.; Hamsten, M.; von Feilitzen, K.; Forsberg, M.; Persson, L.; Johansson, F.; Zwahlen, M.; von Heijne, G.; Nielsen, J.; Pontén, F. Proteomics. Tissue-Based Map of the Human Proteome. Science 2015, 347 (6220), 1260419.

(44) Palma, A.; Tinti, M.; Paoluzi, S.; Santonico, E.; Brandt, B. W.; Hooft van Huijsduijnen, R.; Masch, A.; Heringa, J.; Schutkowski, M.; Castagnoli, L.; Cesareni, G. Both Intrinsic Substrate Preference and Network Context Contribute to Substrate Selection of Classical Tyrosine Phosphatases. J. Biol. Chem. 2017, 292 (12), 4942–4952.

(45) Huang, C.-F.; Mrksich, M. Profiling Protein Tyrosine Phosphatase Specificity with Self-Assembled Monolayers for Matrix-Assisted Laser Desorption/Ionization Mass Spectrometry and Peptide Arrays. ACS Comb. Sci. 2019, 21 (11), 760–769.

(46) Selner, N. G.; Luechapanichkul, R.; Chen, X.; Neel, B. G.; Zhang, Z.-Y.; Knapp, S.; Bell, C. E.; Pei, D. Diverse Levels of Sequence Selectivity and Catalytic Efficiency of Protein-Tyrosine Phosphatases. Biochemistry 2014, 53 (2), 397–412.

(47) Wasserman, W. W.; Sandelin, A. Applied Bioinformatics for the Identification of Regulatory Elements. Nat. Rev. Genet. 2004, 5 (4), 276–287.

(48) Imamura, H.; Wagih, O.; Niinae, T.; Sugiyama, N.; Beltrao, P.; Ishihama, Y. Identifications of Putative PKA Substrates with Quantitative Phosphoproteomics and Primary-Sequence-Based Scoring. J. Proteome Res. 2017, 16 (4), 1825–1830.

(49) Irving, E.; Stoker, A. W. Vanadium Compounds as PTP Inhibitors. Molecules 2017, 22 (12). https://doi.org/10.3390/molecules22122269.

(50) Ostman, A.; Frijhoff, J.; Sandin, A.; Böhmer, F.-D. Regulation of Protein Tyrosine Phosphatases by Reversible Oxidation. J. Biochem. 2011, 150 (4), 345–356.

(51) Londhe, A. D.; Bergeron, A.; Curley, S. M.; Zhang, F.; Rivera, K. D.; Kannan, A.; Coulis, G.; Rizvi, S. H. M.; Kim, S. J.; Pappin, D. J.; Tonks, N. K.; Linhardt, R. J.; Boivin, B. Regulation of PTP1B Activation through Disruption of Redox-Complex Formation. Nat. Chem. Biol. 2020, 16 (2), 122–125.

(52) Lee, S. R.; Kwon, K. S.; Kim, S. R.; Rhee, S. G. Reversible Inactivation of Protein-Tyrosine Phosphatase 1B in A431 Cells Stimulated with Epidermal Growth Factor. J. Biol. Chem. 1998, 273 (25), 15366–15372.

(53) Kalesh, K. A.; Tan, L. P.; Lu, K.; Gao, L.; Wang, J.; Yao, S. Q. Peptide-Based Activity-Based Probes (ABPs) for Target-Specific Profiling of Protein Tyrosine Phosphatases (PTPs). Chem. Commun. 2010, 46 (4), 589–591.

(54) Olsen, J. V.; Blagoev, B.; Gnad, F.; Macek, B.; Kumar, C.; Mortensen, P.; Mann, M. Global, in Vivo, and Site-Specific Phosphorylation Dynamics in Signaling Networks. Cell 2006, 127 (3), 635–648.

(55) Talha, S.; Harel, L. Early Effect of Growth Factors (EGF + Insulin) upon ATP Turnover in 3T3 Cells. Exp. Cell Res. 1983, 149 (2), 471–481.

(56) Emmerson, K.; Roehrig, K. Epidermal Growth Factor (EGF) Stimulation of ATP Citrate Lyase Activity in Isolated Rat Hepatocytes Is Age Dependent. Comp. Biochem. Physiol. B 1992, 103 (3), 663–667.

(57) Makukhin, N.; Ciulli, A. Recent Advances in Synthetic and Medicinal Chemistry of Phosphotyrosine and Phosphonate-Based Phosphotyrosine Analogues. RSC Med Chem 2020, 12 (1), 8–23.

(58) Lemmon, M. A.; Schlessinger, J. Cell Signaling by Receptor Tyrosine Kinases. Cell 2010, 141 (7), 1117–1134.

(59) Barr, A. J.; Ugochukwu, E.; Lee, W. H.; King, O. N. F.; Filippakopoulos, P.; Alfano, I.; Savitsky, P.; Burgess-Brown, N. A.; Müller, S.; Knapp, S. Large-Scale Structural Analysis of the Classical Human Protein Tyrosine Phosphatome. Cell 2009, 136 (2), 352–363.

(60) Coles, C. H.; Shen, Y.; Tenney, A. P.; Siebold, C.; Sutton, G. C.; Lu, W.; Gallagher, J. T.; Jones, E. Y.; Flanagan, J. G.; Aricescu, A. R. Proteoglycan-Specific Molecular Switch for RPTPσ Clustering and Neuronal Extension. Science 2011, 332 (6028), 484–488.

(61) Ostman, A.; Frijhoff, J.; Sandin, A.; Böhmer, F.-D. Regulation of Protein Tyrosine Phosphatases by Reversible Oxidation. J. Biochem. 2011, 150 (4), 345–356.

(62) Fujikawa, A.; Sugawara, H.; Tanga, N.; Ishii, K.; Kuboyama, K.; Uchiyama, S.; Suzuki, R.; Noda, M. A Head-to-Toe Dimerization Has Physiological Relevance for Ligand-Induced Inactivation of Protein Tyrosine Receptor Type Z. J. Biol. Chem. 2019, 294 (41), 14953–14965.

(63) Huang, D. W.; Sherman, B. T.; Lempicki, R. A. Systematic and Integrative Analysis of Large Gene Lists Using DAVID Bioinformatics Resources. Nat. Protoc. 2009, 4 (1), 44–57.

(64) Huang, D. W.; Sherman, B. T.; Lempicki, R. A. Bioinformatics Enrichment Tools: Paths toward the Comprehensive Functional Analysis of Large Gene Lists. Nucleic Acids Res. 2009, 37 (1), 1–13.

(65) Tyanova, S.; Temu, T.; Cox, J. The MaxQuant Computational Platform for Mass Spectrometry-Based Shotgun Proteomics. Nat. Protoc. 2016, 11 (12), 2301–2319.

